# Ecology, molecules and colour: Multivariate species delimitation and conservation of Harlequin poison frogs

**DOI:** 10.1101/050922

**Authors:** Andres Posso-Terranova, Jose A. Andres

## Abstract

We propose a iterative protocol for delimiting species under the generalized lineage concept (GLC) based on the multivariate clustering of morphological, ecological, and genetic data. Our rationale is that the resulting groups should correspond to evolutionarily independent metapopulation lineages because they reflect the common signal of different secondary defining properties (ecological and genetic distinctiveness, morphological diagnosability, etc.), implying the existence of barriers preventing or limiting gene exchange. We applied this method to study a group of highly endangered poison frogs, the *Oophaga histrionica* complex. In our study case, we use next generation targeted amplicon sequencing to obtain a robust genetic dataset that we then combined with patterns of morphological and ecological divergence. Our analyses revealed the existence of at least five different species in the histrionica complex (three of them new to science) occurring in very small isolated populations outside any protected areas. More broadly, our study exemplifies how transcriptome-based reduction of genomic complexity and multivariate statistical techniques can be integrated to successfully identify species and their boundaries.

**In memoriam:** “*I propose that each species has a distinctive life history, which include a series of stages that correspond to some of the named species concepts”*

*Richard G. Harrison*

*1945-2016*

## Introduction

The delimitation of species boundaries is often ambiguous and we continuously debate the taxonomic status of many threatened organisms. In many cases, conflicting species concepts based on operational properties (e.g. reproductive isolation, Mayr 1942; ecological distinctiveness, Van Valen 1976; morphological diagnosability, Helbig et al. 2002; genealogical exclusivity, Baum and Donoghue 1995; etc) have led to confusing situations in which particular populations are considered to be species–or not–under alternative species concepts. For instance, the Florida panther, thought to be a unique (sub)species based on its morphological and ecological distinctiveness (Hedrick 1995) is genetically considered an allopatric population of the North American cougar (*Puma c. cougar;* Culver et al. 2000).

While the adoption by many taxonomists of the General Lineage Concept (GLC) (de Queiroz 2005a, b) has helped to ameliorate some taxonomic controversies, the delimitation of species is still far from trivial. Distinguishing between local adaptation, isolation by distance and species boundaries can be difficult in highly mobile organisms (Brown et al. 2007). Similarly, inter-and intra-specific variation can also be very difficult to differentiate in those taxa characterized by allopatric lineages or low levels of morphological diversity (Barley et al. 2013). As a result, in the last decade we have witnessed the rapid succession of methodological approaches for identifying species boundaries. Thanks to the advent of sophisticated genomics tools and increasing computational power, new methods have been developed to delimit species based exclusively on genetic data (Heled and Drummond 2010; Leaché and Fujita 2010; Huelsenbeck et al. 2011; Grummer et al. 2013; Leaché et al. 2014; Ruane et al. 2014; Zhang et al. 2014; Yang 2015). Similarly, powerful statistical approaches have been proposed to use gaps in morphological variation or ecological discontinuities as the criteria for species delimitation (e.g. Varcarcel and Vargas 2010; Zapata and Jimenez 2012). Despite the widespread use of the above approaches, it is arguable whether species delimitation should be based on just a single line of evidence (Carstens et al. 2013) Instead, an integrative approach should be able to provide the best inferences about species boundaries (Padial and De la Riva 2006; Padial and De La Riva 2010; Padial et al. 2010).

Integrative taxonomy (Dayrat 2005), combines several lines of evidence to support or reject the hypothesis that populations are independently evolving lineages (*i.e.* species; Padial and De la Riba 2009). Under this framework, different lines of evidence can either disagree over the number of species, or agree over the number of species but disagree over the assignment of individuals into species. So, how much agreement must different taxonomic characters show to consider a population (or a group of populations) a separate species? A prevalent strategy has been deriving species hypotheses by consensus, congruence or cumulation (*sensu* Padial et al. 2010) of separately analysed datasets (e.g. Cardoso et al. 2009; Vieites et al. 2009; Dejaco 2012; Yeates et al. 2011). One limitation of these methods is that they cannot extract the common signal that may emerge from the simultaneous analysis of all taxonomic characters. A second limitation is that the resulting species boundaries are likely to be affected by order in which different characters (criteria) are introduced in the analyses. These methodological shortcomings, common in many taxonomic studies, need to be addressed.

One of our main objectives is to provide a total-evidence procedure to analyse morphological, ecological, and genetic variation that can serve to assign specimens into diagnosable species. For this purpose, we performed a comprehensive study of the Harlequin poison frog (*Oophaga histrionica* Myers and Daly 1975) species complex. We have two reasons to focus on this system. First, polytypic taxa present important challenges for species delimitation. Second, specimens of these frogs are actively sought in the wildlife pet trade. Intense illegal trafficking and habitat loss associated with the proliferation of illegal crops have pushed these frogs to near extinction. Thus, it is critically important to delimit conservation units (at or below the species level) that can guide the implementation of the appropriate taxon-based conservation legislature (e.g. CITES trade treaty; Lynch and Arroyo 2009).

Harlequin poison frogs inhabit the lowland Pacific rainforests of the Colombian Choco. Individuals from different populations can either be striped, spotted or relatively homogenous, and their colours range from bright red and yellow to dull green and blue (Figure 1). This phenotypic variability combined with their patchy distribution and strong population structure suggests that *O. histrionica* may, in fact, be a complex of several different species. However, early taxonomic studies (Berthold 1846; Myers and Daly 1976) and preliminary genetic data from geographically scattered individuals (Grant et al. 2006; Brown et al. 2011) have failed to recognized the morphological diversity and potential ecological divergence in this group (Lötters 1992; Lötters et al. 1999; Medina et al. 2013; Posso-Terranova and Andres 2016). To date, only two nominal species are recognized: *O. histrionica* (*sensu lato*) and *O. lehmanni* (Myers and Daly 1976). While the former taxon has a widespread distribution, *O. lehmanni* is a micro-endemic species restricted to a few small submontane wards of the central Choco (Figure 1a).

**Figure 1.**
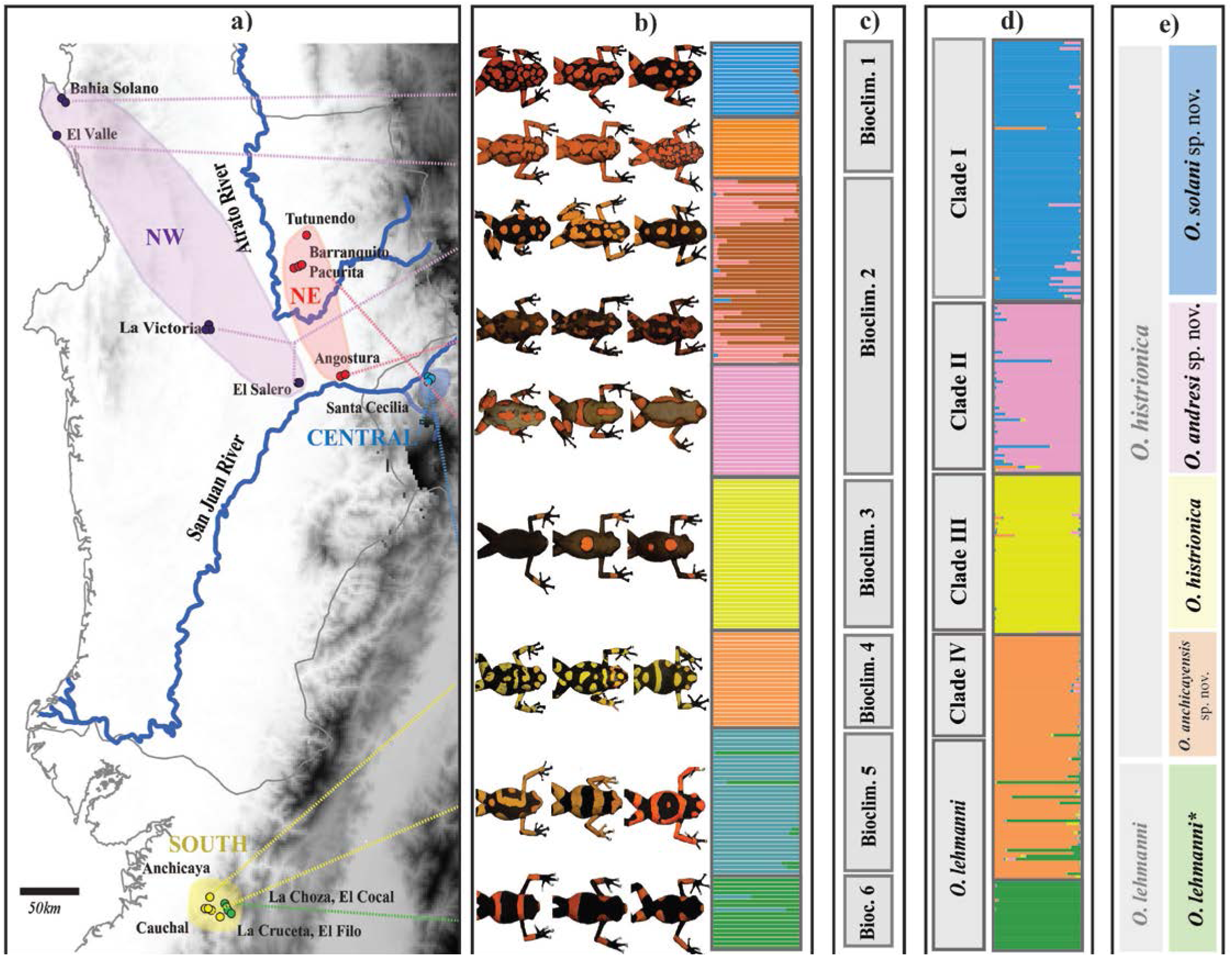
Left to right. a) Map of the sampling localities. b) Representative examples of phenotypic variation found in the “*histrionica*” complex and posterior assignment probabilities to each of the nine distinct morphological groups inferred by the DAPC analysis of 50 morphological variables (see Fig. S1a in Appendix 3 provided in Supplementary Material). c) Schematic representation of the different bioclimatic niches occupied by different populations as inferred by the NMDS analysis of 19 bioclimatic variables and altitude. The multivariate representation of the Gaussian bioclimatic clusters is provided in Figure S2 in Appendix 3 provided in Supplementary Material. d) Grey: Monophyletic mitochondrial clades (I to IV and *O. lehmanni*) as inferred by Bayesian and ML methods. A detailed topology of the phylogenetic tree including *O. sylvatica* and *O. occultator* as outgroups is shown in Figure S3 in Appendix 3 provided in Supplementary Material. The corresponding parsimony haplotype network is available in Figure S4 in Appendix 3 provided in Supplementary Material. Color: Posterior probability of assignment to five genetic clusters based on Bayesian analysis of variation at microsatellite and transcriptome-derived haplotypes. Graphical representation of the corresponding PCoAs are provided in Figures S5a and S5b in Appendix 3 in Supplementary Material. e) In grey, species delimitation based on Myers and Daly (1976) and Funkhouser (1956). Coloured-boxes represent the new delimitation resulting from our multivariate integrative analyses. *sp. nov*. indicate the new species proposed, and described, in this paper.

In this study, we used next generation sequencing (NGS) to obtain haplotypes that are then analysed using multivariate clustering approaches. This approach, similar to the species delimiting method of Hausdorf and Hennig (2010) and the species assignment method of Edwards and Knowles (2014), combines transcriptome polymorphisms with mtDNA and microsatellites variation to generate a comprehensive genetic dataset that is then combined with patterns of morphological and ecological divergence. The resulting common signal is used to establish species delimitations. To evaluate the power of our approach to identify independent evolving lineages (i.e. species), we compared its results to those obtained by other “traditional” integrative methods (see Edwards and Knowles 2014). Our analyses revealed the existence of at least five different species in the *histrionica* complex (three of them new to science) occurring in very small isolated populations outside any protected areas. More broadly, our study exemplifies how targeted amplicon sequencing and multivariate statistical techniques can be integrated to successfully identify species and their boundaries.

## Materials and Methods

### Specimen sampling

We sampled 310 individuals in 15 different geographic locations across the known geographic distribution of the *O. histrionica* complex (Myers and Daly 1976; Lötters 1992; Lötters et al. 1999) (Fig. 1a). For each of the sampled individuals, we recorded its geographic location using a handheld GPS unit. High-resolution images and tissue samples were then collected as follows: raw digital pictures (Nikkon Electronic Format: NEF) were taken using a Nikkon D610 digital camera against a Spectralon grey reflectance standard (Labsphere, Congleton, UK). Images were then linearized with respect to light intensity (Stevens et al. 2007), and stored as TIFF images (lossless compression) to preserve fidelity for further analysis. Tissue samples were obtained by toe-clipping (Phillott et al. 2011). Toe-clips were conserved in 95% ethanol until DNA extraction. Genomic DNA was extracted using the MasterPure^TM^ DNA purification kit (Epicentre, Madison, WI, USA) following the manufacturer’s recommendations.

All samples were obtained in compliance with the Ethic Committee of the Universidad Nacional de Colombia (sesión del 22 de Julio, Acta No.03, July 22^nd^ 2015), under the research permits granted by Autoridad Nacional de Licencias Ambientales ANLA, Resolución 0255 del 12 de Marzo de 2014 and Resolución 1482 del 20 de Noviembre de 2015.

### Morphological clustering

Species are commonly delimited on the basis of gaps in patterns of morphological variation (Zapata and Jiménez 2012). Hence, we estimated the number of distinct morphological clusters (species) using hierarchical and discriminant grouping methods. Dorsal and ventral pictures of each specimen were scaled using ImageJ v. 1.48 (Sheffield 2007) and a known metric reference. For each specimen, we then measured a total of 49 variables (n_dorsal_= 25; n_ventral_= 24) related to morphology, pattern of coloration and coloration intensity (Table S1 in Appendix 1). All variables were standardized prior to subsequent analyses. First, we performed a hierarchical cluster analysis as implemented in PAST v. 3.08 with bootstrap-and PERMANOVA-based significance tests (Hammer et al. 2001). In addition, we also performed a discriminant analysis of principal components (DAPC) as implemented in the R package *Adegenet* v 2.0.0. (Jombart et al. 2010). This method has the advantage that allows for a probabilistic assignment of specimens to each group. To determine the optimal number of morphological clusters (Km), we ran the sequential *K*-means (*find.clusters, K*=1-19) algorithm and identified one showing the lowest Bayesian Information Criterion (BIC) (Jombart et al. 2010). To avoid overfitting, the individual assignment of specimens was based on 11 axes (*optim.a.score*) representing 86% of the variation in the dataset.

### Ecological clustering

To identify ecological discontinuities and determine the number of environmental clusters (Ke) we performed a non-metric multi-dimensional scaling (NMDS) analysis as proposed by Edwards and Knowles (2014). Briefly, for each sampled individual, we extracted high resolution (30 arc-seconds) environmental data from 19 bioclimatic variables (www.worldclim.org) along with altitude data (STRM90 http://srtm.csi.cgiar.org/) (Hijmans et al. 2005). Bioclimatic data were standardized and reduced to PC-scores along 4 axes (explaining > 90% of the variation) using the *dudi.pca* function in the Vegan R-package (Oksanen et al. 2013). Next, geographic distances among sampled specimens (function: *Distance*, Diva-Gis v. 7.5; http://www.diva-gis.org/) (Hijmans et al. 2001) were converted to continuous variables (function: *pcnm*. Vegan R-package) and combined with the bioclimatic scores before generating the final Gower-distance environmental dataset. Then, we used non-metric multi-dimensional scaling (NMDS) and one-way PERMANOVA to estimate Ke.

### Genetic clustering

*Mitochondrial DNA (mtDNA):* Mitochondrial genetic subdivision is often considered a leading criterion of interspecific differentiation. To determine the number of mitochondrial clusters (Kmt), we applied phylogenetic and multivariate procedures. For a subsample of 107 individuals (including *O. sylvatica* (n=6) and *O. occultator* (n=4) as “*bona fide*” control species) we amplified and Sanger-sequenced a 523 bp of the COI gene (see Hauswaldt et al. 2011; Brusa et al. 2013; Medina et al. 2013 for detailed molecular protocols). Aligned sequences (*Clustal-W)* (Thompson et al. 1994) were used to estimate the model of nucleotide substitution using the corrected Akaike Information Criterion as implemented in jModelTest v2.1.4 (Posada 2008). MtDNA phylogenetic trees were constructed under different Bayesian and maximum likelihood optimality criteria and a TCS haplotype network was also plotted (Appendix 2). Individual pairwise genetic distances were calculated using MEGA6 (Tamura et al. 2013), and a Principal Coordinate analysis (PCoA) was carried out upon the similarity matrix (Reeves and Richards 2011). Finally, to delimit independent evolving linages we also used a generalized mixed-Yule-coalescent (GMYC) method capable of fitting within-and between-species branching models (Zhang et al. 2013) (Appendix 2).

*Microsatellites and transcriptome-derived haplotypes*: Twelve microsatellite loci were isolated from a di-, tri-and tetra-mer enriched library constructed from a single *O. histrionica* specimen using pyrosequencing technology (454GS-FLX. Roche Applied Science) as described in Andres and Bogdanowicz (2011) (Table S2 in Appendix 1). All individuals (n= 310, nlocalities= 15) were genotyped using multiplex-PCR and fluorescently-labelled primers as described in Appendix 2,.

To further identify polymorphic nuclear loci for multiplex PCR we used a transcriptome-based approach. *De novo* skin transcriptome sequencing and assembly Table S3 in Appendix 1) from a single *O. lehmanni* individual was carried out as described in Appendix 2. The resulting transcriptome was utilized to design primers for 50 nuclear markers (250-350 bp), each of them representing a single randomly selected unigene. To reduce ascertainment bias, we selected eight individuals from a broad geographic range to screen each locus for robust PCR using genomic DNA as template. After stringent selection, 20 remaining loci were used to genotype all 310 specimens in two multiplex PCR groups using the QIAGEN Multiplex PCR Kit. Individual specimens and loci were barcoded using Illumina^TM^ S5 and N7 Nextera primers following the manufacturer’s conditions. Amplicons were paired-end sequenced (2X 250 bp) on a Illumina MiSeq at the Cornell University’s BioResource Center.

Post-sequencing data processing required extracting reads from the Miseq run and assigning them to the appropriate specimen and locus with a custom *Perl* script that discarded low quality reads, trimmed adapters, overlapped paired-reads, identified reads corresponding to each locus, collapsed identical reads for each individual, and identified the two most frequent haplotypes for each individual at all loci (quality score: Q12, minimum overlap: 20, mismatch rate: 0.05). At any given locus, an individual was scored as heterozygous if the minor allele frequency was ≥ 20%. Individuals that fail to amplify in more than 80% of the loci were excluded from the dataset, and any sample by locus cells with fewer than 10 total reads were considered missing data.

The number of genetic clusters based on microsatellite loci (Ksat), transcriptome-derived haplotypes (Ktrn), and both datasets combined (Knuc) was estimated using three allele frequency-based methods (STRUCTURE, STRUCTURAMA, and PCoA). While the first two methods estimate the number of clusters by minimizing the deviations from Hardy-Weinberg equilibrium, the latter one finds clusters based on the Nei’s distance (Pritchard et al. 2000; Hubisz et al. 2009; Peakall and Smouse 2012). Detailed information on the different models, parameters and run conditions is described in Appendix 2.

### Total-evidence species delimitation

The multivariate species delimitation approach described in this paper assumes the GLC species concept, and is based on the conceptual and methodological frameworks of Hausdorf and Henning (2010) and Edwards and Knowles (2014). First, a provisional species hypothesis is formulated based on a small balanced geographical sampling of the focal system and the assumption that species can be defined as distinguishable groups (i.e. multivariate clusters) of individuals with similar phenotypic, genotypic and ecological characteristics (see Discussion). Second, we weighted the strength of the evidence supporting the proposed species hypothesis by evaluating our ability to detect and assign new specimens to the provisional species. Because this multivariate approach does not necessarily take into account historical patterns of diversification, we compared our results to those obtained by using a Generalized Mixed Yule Coalescent (GMYC) model capable of fitting within-and between-species branching models (Zhang et al. 2013).

Accordingly, for a first sample of individuals (n=86, 4-12 ind./population), we generated a new set of standardized orthogonal variables containing the same information as the original datasets using multivariate methods (morphology: PCA, environmental and genetic: PCoA). We then calculated distance matrices for each datatype (morphology and environmental: Gower-distances; mtDNA: TN93-distance; nuclear loci: Nei’s distance) and standardized them using NMDS (*isoMDS*, MASS R-package). To determine the provisional number of species we followed Edwards and Knowles (2014) and retained four dimensions for each data-type and determined the number of Gaussian clusters present in the subsample using the R-package *mclust* and the Bayesian Information Criterion (Fraley and Raftery 2002). Additionally, we also performed hierarchical clustering (average method) with 1000 bootstrap repetitions as implemented in PAST v. 3.08 (Hammer et al. 2001).

To assess the strength of our provisional species delimitation hypothesis, we conducted an identical series of analyses on an independent, much larger dataset (n= 214 individuals) and assessed our ability to recover the same clusters (species) with high individual assignment probabilities. All stress values for NMDS analyses were below 10%, with an absolute maximum of 6%.

### Nomenclature acts

All new species names resulting from this study conform to the requirements of the amended International Code of Zoological Nomenclature (ICZN) and they have been registered in ZooBank (http://Zoobank.org/). All Relevant information as well as the Life Science Identifiers (LSIDs) can be found under the account <u>DACAA96D-2E69-4718-83DD-EB907A83BC95</u>.

## Results

### Morphological and environmental clustering

The morphological hierarchical clustering applied over individuals of the *O. histrionica* complex revealed the presence of nine clearly defined morphological clusters with high node support (>82%; PERMANOVA Bonferroni-adjusted P<0.003; Fig. 1b and Fig. S1 in Appendix 3). The optimal number of morphologic clusters (Km) as defined by DAPC (BIC) was also nine. As a result, both clustering methods were highly congruent in their individual assignments (Fig. 1a). Posterior assignment probabilities (PP) for Km=9 indicated that 93% of individuals were correctly assigned to their morphologic clusters (PP>0.9); 4.7% were considered as ambiguously assigned (PP<0.9) and only 2.3% were incorrectly assigned (*i.e.* assigned to a different cluster with PP>0.9).

Gaussian clustering based on a NMDS analysis of bioclimatic distances partitioned the sampled geographic range of the *O. histrionica* complex into six distinct ecological clusters (KE= 6, stress=0.04) with statistical significance after Bonferroni correction (P<0.0001; Fig. 1c): while *O. histrionica* (*sensu lato*) encompasses four potential bioclimatic niches, putative *O. lehmanni* specimens form two different clusters that correspond to two different morphological clusters (Fig. 1c and Fig. S2 in Appendix 3).

### Genetic clustering

Bayesian (BI) and Maximum Likelihood (ML) COI gene genealogies showed identical topologies and revealed the presence of seven distinct mtDNA clades representing *O. sylvatica*, *O. occultator*, *O. lehmanni* (defined by phenotype and sampling location), and 4 monophyletic, geographically structured clades (I-IV) within *O. histrionica* (*sensu lato*, Fig. 1d and Fig. S3 in Appendix 3): Clade I (NW-populations), Clade II (NE-populations), Clade III (Central populations), and Clade IV (South populations)–See also haplotype network provided in Figure S4 in Appendix 3 –. As expected, on average, the *p-*distances between the “*bona fide* species” (*O. occultator* and *O. sylvatica*), and the *O. histrionica* complex clades were greater (range: 4-6%) than those observed among the four *histrionica* clades (range: 2.2%-2.9%). The single-locus species delimitation analysis by GMYC strongly supported the hypothesis that each of these mtDNA clades represent an independent species (PS>0.96 in all cases).

Clustering analyses of the *O. histrionica* complex based on the two nuclear datasets yield consistent partitions that, in turn, were very similar to that based on mtDNA. Although the PCoA results were mostly concordant, the microsatellite dataset supported a five-cluster partition (Ksat= 5, Fig. S5a in Appendix 3) while transcriptome-derived dataset supported six clusters (Ktrn= 6, Fig. S5b in Appendix 3). The source of discordance is the adscription of the individuals from the localities of La Choza and El Cocal. The analysis of the transcriptome-derived haplotypes suggested that these populations represent an intermediate cluster between the Southern populations of *O. histrionica* (*sensu lato*)-Anchicaya and El Cauchal-and the *O. lehmanni* populations of La Cruceta and El Filo (Fig. 1a). In contrast, the microsatellite-based analysis clustered these individuals with either the Southern *O. histrionica* or the *O. lehmanni* cluster. These results are consistent with La Choza and El Cocal being admixed populations. As expected under this admixture hypothesis, La Choza-Cocal specimens have intermediate phenotypes between those of *O. lehmanni* and Southern *O. histrionica* (Fig. 1b), and either Clade IV or *O. lehmanni* mtDNA haplotypes (Fig. 1d).

Bayesian clustering (STRUCTURE) analyses for the combined data set (microsatellites + transcriptome haplotypes) showed that the estimated log probability *[Ln P(D)*] plateaued at KNUC = 6 with the second-order rate of change of *K* (*ΔK*) reaching its maximum at KNUC = 5, suggested the existence of five (potentially six) distinct genetic clusters. Assuming KNUC = 5, the first cluster grouped all *O. lehmanni* specimens, while the other four clusters corresponded to the mtDNA clades I-IV (Fig. 1d). There were 37 individuals exhibiting admixed ancestry *(q =* 0.1–0.9), and 73% of them were found in four populations (Angostura and El Salero n= 14, Clade I *x* Clade II; La Choza and El Cocal, n= 13, Clade IV *x O. lehmanni)*. The geographic location of these admixed populations (Figure 1a) suggests that these localities represent two different contact zones, one between the NW and the NE demes, and the second one between the South and the *O. lehmanni* ones. The results of the STRUCTURAMA analyses also supported the partition of KNUC= 5 with a posterior probability of 74% (Table 1).

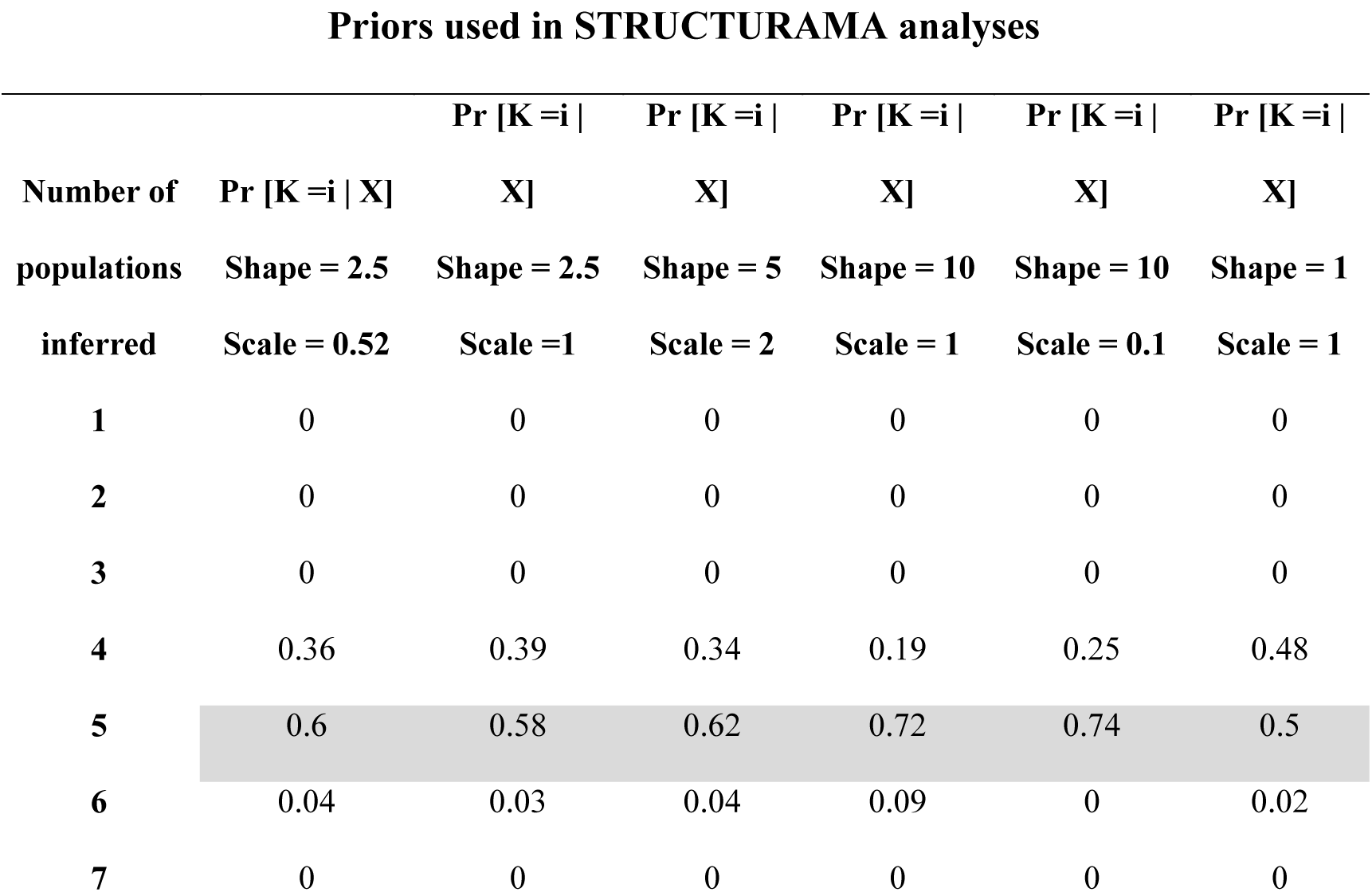

Table 1. Results from STRUCTURAMA analyses at a range of priors for shape and scale using the *O. histrionica* species complex genetic dataset from microsatellite (n=6) and transcriptome-based markers (n=13). Optimal number of inferred populations is shown in light grey.

### Species delimitation by integrative analysis

To first delimit the number of species, we used an integrated dataset composed of 12 multivariate dimensions (see Methods) and two different clustering methods: first, we used a model-based Gaussian clustering. The best model corresponded to an equal-covariance and diagonal distribution model (VEI) (Fraley and Raftery 2002) with the BIC plateaued at K= 5 (Fig. S6 in Appendix 3). Bootstrap likelihood ratio tests (*MclustBootstrap*, n*= 999*) for K = 1-7 supported the existence of five lineages representing *O. lehmanni* (including the genetically admixed populations of La Choza – El Cocal), and the S, Central, NE, and NW lineages *of “histrionica*”. Such taxonomic partitioning resulted in 100% assigned individuals (PP ≥ 0.95). The second-best model (K=6) split *O. lehmanni* into ‘pure’ (El Filo–La Cruceta) and admixed (La Choza–El Cocal) populations. Secondly, our hierarchical clustering approach was fully concordant with the Gaussian approach and showed five highly supported clusters (BS>95%) with identical composition of specimens (Fig. 2a). Given these results we subdivided the *histrionica* complex into five provisional species. To test the robustness of these provisional species we performed a second round of clustering analyses in an independent set of specimens (n= 214) and K= 5. Under such delimitation, all individuals were correctly assigned to the expected lineage with PP ≥ 0.95 (Fig. 2b). Clustering analyses excluding mtDNA yield identical assignment results.

**Figure 2.**
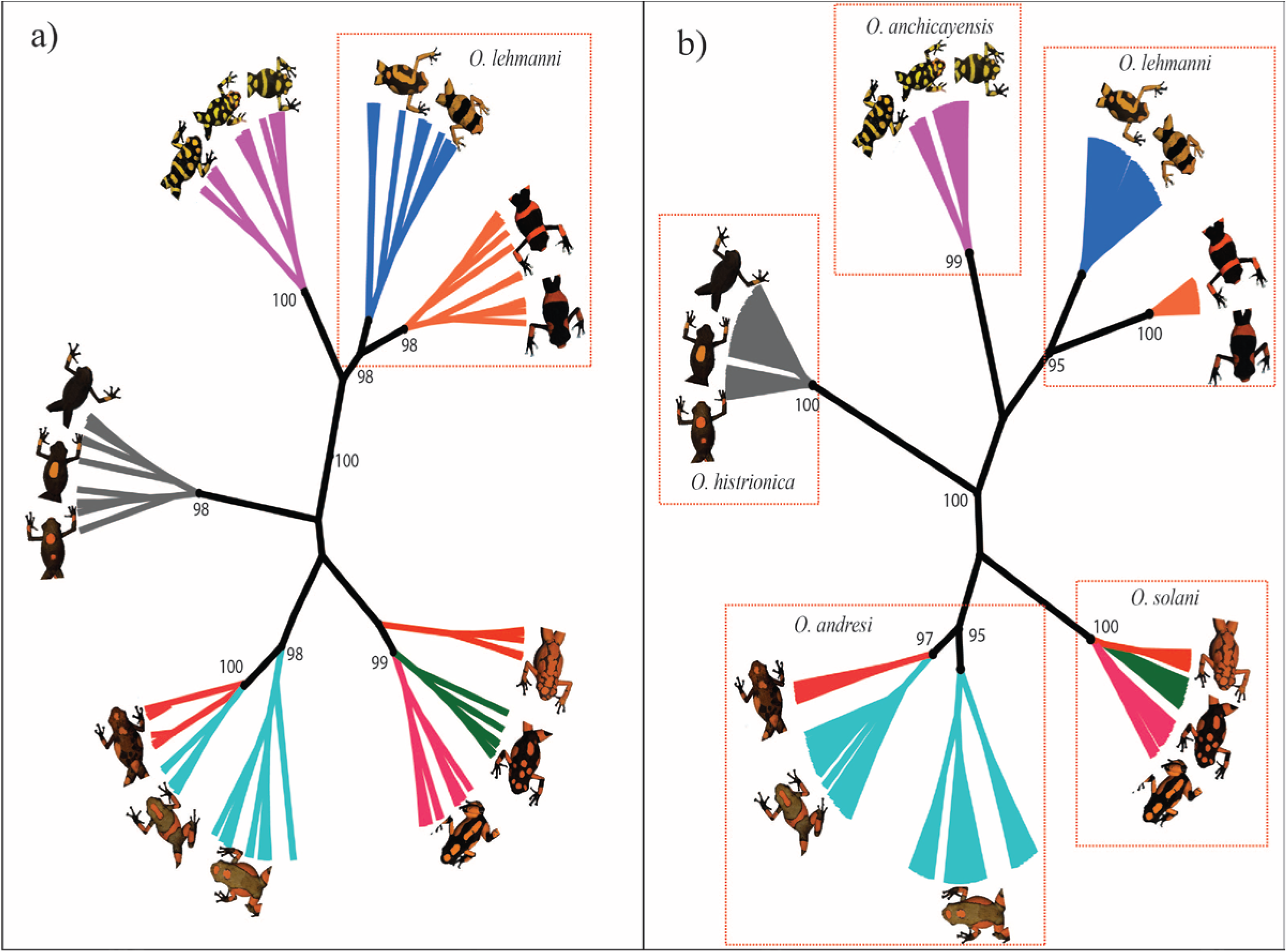
Integrative species delimitation based on the hierarchical clustering standardized orthogonal variables (genetic, ecological and morphological). a) Provisional delimitation resulting from the analysis of 84 specimens. b) Extended clustering analysis of 214 individuals. The colored boxes enclose individual specimens belonging to the same Gaussian cluster, PP ≥ 0.95). Only nodes with support >95% are indicated.

## Discussion

### Conceptual basis

Edwards and Knowles (2014) developed a statistical approach to evaluate hypotheses of species limits based on multivariate analyses and Gaussian clustering of total evidence (morphological, ecological and genetic) datasets. Here, we extend their approach and propose that the iterative implementation of such methodological framework can be used to delimit species under the GLC without any *a priori* species boundaries. Our rationale here is that these multivariate clusters should correspond to evolutionarily independent metapopulation lineages because they reflect the common signal of different secondary defining properties (ecological and genetic distinctiveness, morphological diagnosability, etc.), implying the existence of barriers preventing or limiting gene exchange. It is important to recognize that because of the hierarchical nature of evolution, we expect clustering to occur at many levels. That is, we should observe a continuum of increasing degree of cluster overlap as the levels of genetic, morphological and ecological divergence decreases between species to variation across populations. Thus, depending on the focal system, the inferred clusters might represent inter-population variation, local host races, subspecies, species, etc. So, where along the divergence continuum separate lineages (clusters) should be recognized as distinct species?

To try to answer this question, one may rely on “arbitrary” statistical thresholds. For example, it has been argued that 70% correct multivariate assignment is a reliable threshold for species delimitation in complex groups (Valcárcel and Vargas 2010). However, as noted by Coyne and Orr (2004, pg. 449) species do not have a greater “objective” reality than lower or higher taxa and we do not see a fundamental distinction between species and any other grouping level. A better alternative is to contrast the multivariate clusters with those resulting from the application of multispecies coalescent methods that infer species status from a genealogical perspective.

In the case of the *histrionica* complex studied here, the GMYC analysis showed strong correspondence between the inferred multivariate clusters and the branching pattern of the mtDNA species tree. The only source of discordance was that the integrative analysis grouped a few *O. lehmanni* specimens together with individuals from the ‘*histrionica*’ mtDNA cluster IV (Fig. S3 and S4 in Appendix 3). Multivariate and Bayesian analyses showed that all ‘misclassified’ individuals are of admixed origin (Fig. 1b and 1d), a result that is concordant with previous studies suggesting hybridization between *O. histrionica* (*sensu lato*) and *O. lehmanni* (Medina et al. 2013).

Despite its limitations, we believe that this approach has some practical advantages over other delimitation methods often reliant on genetic data. First, assuming that there is enough variation, this method provides a combination of intrinsic characters (attributes) that can be used to recognize and describe the newly discovered species. Second, because this approach relies in the cumulative effect of independent characters it is likely to be useful to unravel species in early differentiation stages were varying levels of differentiation across genetic, phenotypic and ecological axes are expected (Padial and De La Riva 2010; Padial et al. 2010). Third, this method can be useful for analysing large datasets. Most coalescent based delimitation methods present serious computational challenges when analysing significant numbers of loci, putative species and/or individuals. In contrast, Gaussian clustering requires little computational power and can be easily scalable to include numerous genetic loci as well as large numbers of specimens.

### Taxonomic implications

A key question in our study was to determine whether the extensive phenotypic diversity within the *O. histrionica* complex reflects the existence of several independent taxonomic units *(i.e*. species) or just one (possibly two) highly polytypic species (Myers and Daly 1976; Grant et al. 2006). The application of our integrative multivariate procedure confirmed the taxonomic status of *O. lehmanni* and provided statistical support for the existence of five distinct species of *Oophaga*, which we propose should be formally recognized.

Though the specific status of *O. lehmanni* has casted doubt among taxonomist during the last decades (Lötters 1992; Lötters et al. 1999; Medina et al. 2013), analyses indicated a considerable amount of genetic, ecological and morphological differentiation between the two current nominal species *O. lehmanni* and *O. histrionica* (Myers and Daly, 1976; Fig. 1e). Thus, we recommend maintaining the taxonomic status of *O. lehmanni*. Our study also provides further support for existence of a geographically isolated hybrid lineage (i.e. cluster) of *O. lehmanni* (Medina et al. 2013). Overall, the pattern of genetic additivity of parental marker alleles (Fig. 1d), along with ecological (Fig. 1c) and morphological separation (Fig. 1b) suggests that hybridization between these two lineages may have resulted in the successful establishment of an ecologically distinct lineage. Further genetic studies are required to test this hypothesis.

We also propose the existence of four species within *O. histrionica* (*sensu lato*). These species correspond to each of the four ‘*histrionica’* clusters revealed by our integrative analyses.

Here, we provide the proposed names and descriptions of the newly delimited species. Although detailed morphological descriptions have been previously proposed (Berthold 1846; Silverstone 1975; Myers and Daly 1976; Daly et al. 1978; Lötters 1992; Lötters et al. 1999; Grant et al. 2006), we have included a summarized description of each species with respect to body measurements and a synthesized description of coloration patterns based on our multivariate morphologic analysis. Distributional ranges are based on our own sampling.

We would like to highlight that if we accept species as evolutionary units, we cannot expect or demand immutable species hypothesis, but only the most corroborated ones (Padial and De La Riva 2010). Several lines of evidence and different methodological approaches support the species described in this paper. Therefore, they can be regarded as having high potential of further corroboration. However, the inclusion of other lines of evidence (e.g. acoustic, behavioral, etc.) over an extended geography may result in new, revised hypotheses.

#### Oophaga solani sp. nov. [Mitochondrial clade I]

We designate this species as *O. solani* sp. nov. (syn. *O. histrionica*) based on the mtDNA (PS=1) and nDNA species delimitation support and the general conservative agreement of the integrative analysis (BS=99%). We named this new species after the municipality of Bahia Solano, department of Choco, Colombia where we started part of our field work and found the first populations of this species. Museum specimens AMNH 86979-86985 correspond to this species (Myers and Daly 1976).

*Range*: Based on our own sampling plus data collected from different authors (Silverstone 1975; Myers and Daly 1976; Mendez-Narvaez and Amezquita 2014), this species inhabits the coastal lowlands at the west banks of the San Juan and Atrato rivers which, due their width, presumably represent a major barrier to gene flow between the eastern and western *Oophaga* species (Figure 1a). Populations of this species are distributed in a relatively wider geographic extension of dense rainforest with known populations from Mecana, Bahia Solano (Lat. 6.29640, Lon.-77.37869) and Serranía del Baudo (Silverstone 1973) throughout La Victoria, (Lat. 5.51074, Lon. −76.86979), El Salero, (Lat. 5.36412, Lon.-76.64629), Nuqui, (Lat.5.70652, Lon.-77.26079) until Quebrada Docordo, department of Choco (Lat. 4.56265, Lon.-77.01399). In contrast to *O. histrionica*, this species inhabits lower altitudes (0 to 324 m.a.s.l) with an annual mean temperature of 25.9 °C and annual mean precipitation of 6279 mm.

*Diagnosis*: Body size in these frogs ranges between 53-33 mm (36.2 ±1.7 mm, n=75). This species represents three different morphologic clusters (morphotypes) detected in this study with high bootstrap support (BS>82%) (Fig. S3 in Appendix 3). Individual frogs are black with bright orange or reddish orange spots that are highly variable in size and number (n=3-30); however, the coloured dorsal area remains relatively constant (40-70% of total dorsal area). Colour variation is exemplified by Myers and Daly (1976) (Fig. 10, pp. 209).

#### Oophaga andresi sp. nov. [Mitochondrial clade II]

We designate this species as *O. andresi* sp. nov. (syn. *O. histrionica*) based on the genetic species delimitation (mtDNA and nDNA, PS=1) and the general conservative agreement of the integrative analysis (BS=98%). We named this new species after our beloved friend Andres Orrego-Ortiz, who passed away at a very young age in Palmira, Valle del Cauca, Colombia, while we were still collecting and analysing data there. Museum specimens representing this species are AMNH 85186-85189 and AMNH 86957-86960 (Myers and Daly 1976)

*Range*: This species is distributed at the east banks of the Atrato river which probably represent a geographic barrier between *O. solani* and *O. andresi* (Fig. 1a). After confirmation of the presence of this species in those geographic locations originally visited by Myers and Daly (1976) plus our own field records, we consider that populations of *O. andresi* are distributed from the surrounding areas of Tutunendo, Choco (Lat.5.75219, Lon.-76.53634) throughout Pacurita, Choco (Lat 5.646565, Lon.-76.56655), Bagado, Choco (Lat.5.38121, Lon.-76.42759) until Vereda La Angostura, Playa de Oro, Choco (Lat. 5.32194, Lon.-76.43090). Similar to *O. solani*, this species inhabits the relatively warmer lowlands (annual mean temperature of 26.6 °C; altitude 55 to 168 m.a.s.l). However, *O. andresi* exhibits the higher requirements of humidity, inhabiting locations with annual mean precipitation >7500 mm.

*Diagnosis*: Frogs of this species are similar in body size to *O. histrionica* (34.9 ±2.4 mm, n=43) and are represented by two distinct morphotypes detected in this study (BS>91%) (Fig. S1 in Appendix 3). The basic and most common coloration pattern is represented by one to three large orange-reddish-yellow spots and bracelets on a background ranging from light to dark brown, but never black in colour. Detailed description of variation in spot sizes and distribution, as well as a representation of morphologic variation is presented in Myers and Daly (1976) as “*Playa de Oro*” frogs (Fig. 8-9, pp.206-208).

#### Oophaga histrionica Berthold 1846. [Mitochondrial clade III]

*Oophaga histrionica* (syn. *Dendrobates histrionicus*) was originally described as a species (Berthold 1846) based on descriptions of an unknown holotype (reproduced in Fig. 6, pp. 200, Myers & Daly, 1976) phenotypically similar to frogs occurring on the upper Rio San Juan, in the Department of Risaralda, Colombia. In this study, *O. histrionica* was highly supported as a unique independent evolutionary lineage by all individual lines of evidence and the integrative analysis (Fig. 1e). A detailed description of coloration and pattern variation of this species is presented in Myers & Daly (1976) as “*Santa Cecilia*” frogs (pp. 205). Museum specimens that represent this species are deposited at the American Museum of Natural History (AMNH) under collection numbers AMNH 85159-85161 (Myers and Daly 1976).

*Range*: Based on our evidence, and contrary to the previously described range, this species is distributed in a narrow geographic range at an altitude between 342-410 m (mean 394 m). Populations of this species are geographically restricted to less than 30 km^2^ in the surrounding areas of the municipality of Santa Cecilia, Colombia (Lat. 5.30336, Lon. −76.21662) with annual mean temperature of 25.6 °C and annual mean precipitation of 5267 mm.

*Diagnosis:* Frogs of this species are significantly smaller in body size than other species of the complex (32.9 ±1.7 mm, n=28). This species represent a clearly defined and unique morphologic cluster (BS=82%) (Fig. S1 in Appendix 3) where individual frogs have light brown to black bodies with usually one or two large orange or red-orange spots on the back. In rare cases, the dorsal marking is absent or difficult to notice. Morphological variation is shown in Myers and Daly (1976) (Fig. 7, pp. 205).

#### Oophaga anchicayensis sp. nov. [Mitochondrial clade IV]

We designate this species as *O. anchicayensis* sp. nov. (syn. *O. histrionica*) based on the genetic species delimitation (mtDNA and nDNA, PS=1) and the general conservative agreement of the integrative analysis (BS=100%). We named this new species after the type locality Anchicaya, Buenaventura, Colombia. Museum specimens representing this species are deposited at the Universidad del Valle, Cali (UVC 12431-52) (Lötters et al. 1999), at the Institut Royal des Sciences Naturelles de Belgique (IRSNB) (MRHN 1038) and AMNH 10613-15 (Silverstone 1975).

*Range:* This species is located at southern localities (Fig. 1a) and its geographic range correspond to <170 km^2^ at Valle del Cauca department, Colombia, at higher elevations when compared with other species of the genus (616-791 m.a.s.l). Populations of this species has been reported at Delfina, Buenaventura (Lat.3.81364, Lon.-76.81305), Cisneros (Lat.3.79955, Lon.-76.80305), Rio Zabaletas, Buenaventura (Lat.3.72003, Lon.-76.91802), Bajo Anchicaya, Dagua (Lat.3.63621, Lon.-76.94299), El Danubio, Dagua (Lat.3.62210, Lon.-76.90199) and vereda El Cauchal, Dagua (Lat.3.61712, Lon.-76.865). In contrast to *O. solani* and *O. andresi*, this species inhabits cooler and dryer geographic regions (annual mean temperature=24.4 °C; annual mean precipitation=2419 mm).

*Diagnosis*: These frogs are significantly bigger in body size than other species of the complex (41.2 ±1.3 mm, n=29). Background coloration is always completely black with dorsal yellow, red-orange or orange spots highly variable in number (3 to 20) and size. Dorsal pattern sometimes includes median bracelets as *O. lehmanni* (Myers and Daly 1976) that are incompletely formed ventrally in most cases. Some populations as “*Delfina*” and “*Cisneros*” (Silverstone 1975; Medina et al. 2013) present an increased reddish coloration towards the head and mouth (first frontal quarter of size length). This pattern is locally known as “*red head*” frogs and is shown in Lötters et al. (1999) (Fig. 3, pp. 27) However, this species is represented by a single morphological cluster with high statistical support (BS=95%) (Fig. S1 in Appendix 3).

### Conservation implications

This paper also illustrates how understanding lineage diversification can inform wildlife management and has relevance to the conservation efforts of South American poison frogs. The new integrative taxonomy proposed here has very important conservation implications as it revealed that some of the species should be considered amongst the most vulnerable species of Neotropical frogs: of the five species (*O. histrionica*, *O. lehmanni*, *O. solani*, *O. andresi*, and *O. anchicayensis*) two of them (*O. histrionica* and *O. lehmanni*) are single-locality endemics and should be both considered critically endangered. The inclusion of *O. lehmanni* as a critically endangered species more than 10 years ago led to the proposal of several conservation programs and regional legal policies aiming to preserve the remaining populations and their habitat (Bolivar et al. 2004; Valencia-Zuleta et al. 2014). Our study reveals that the distribution of *O. histrionica* is restricted to a very small area (< 30 km^2^), and the only 3 known populations of this microendemic species should probably be subject of similar actions. At the very least, *O. histrionica* should be listed as Critically Endangered in the UICN list and the resolution N^0^ 0192-2014 (Ministerio de Ambiente y Desarrollo Sostenible) that establishes the Colombian list of endangered species. Also, given that these two species are highly sought-after in the pet trade market, it will be desirable to include them in the (Appendix I) of the CITES treaty to prevent the commercial trade of wild-caught specimens.

The conservation status of the newly described species (*O. anchicayensis*, *O. andresi* and *O. solani*) is less clear as they all have relatively wide, discontinuous, distributions. Although their effective population sizes are likely to be large, preliminary data suggest that populations are strongly structured. Therefore, several management (MUs) and evolutionary significant (ESUs) units (Funk and Richardson 2002) should probably be considered. In the absence of any better information regarding species population ecology, density or abundance, we consider that these species should be temporarily categorized as vulnerable to extinction. Regardless of their conservation status none of the *Oophaga* species described in this paper occur in protected areas. Ideally, once included in the appropriate endangered-species list, these charismatic frogs will serve as umbrella species for the future conservation of entire ecosystems.

## Acknowledgements

We dedicate this paper to the memory of Richard G. Harrison, beloved friend and inspirational mentor. We would like to thank the members of his lab at Cornell University for their comments on an early version of this manuscript and Biologists Pablo Palacios and Ramon Ramirez for their assistance during the initial field work.

## APPENDIX 1: Supplementary Tables

**Table S1:**
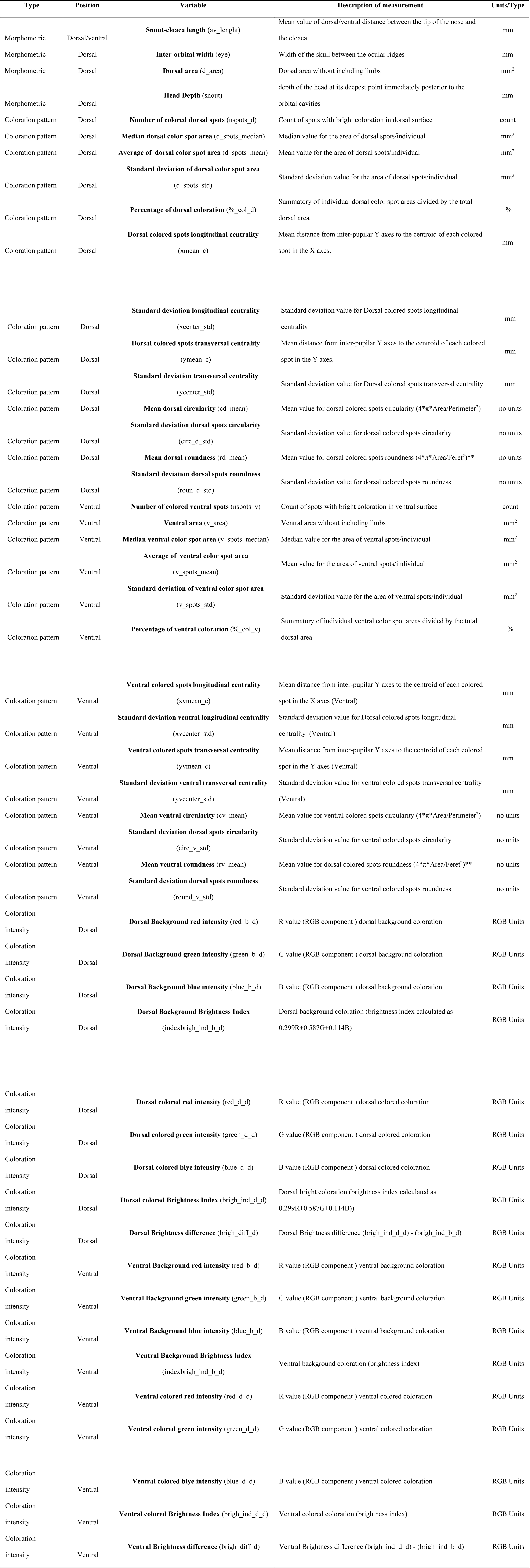
Description of 49 morphologic variables used for the estimation of morphological clusters and combinatory species delimitation.

**Table S2.**
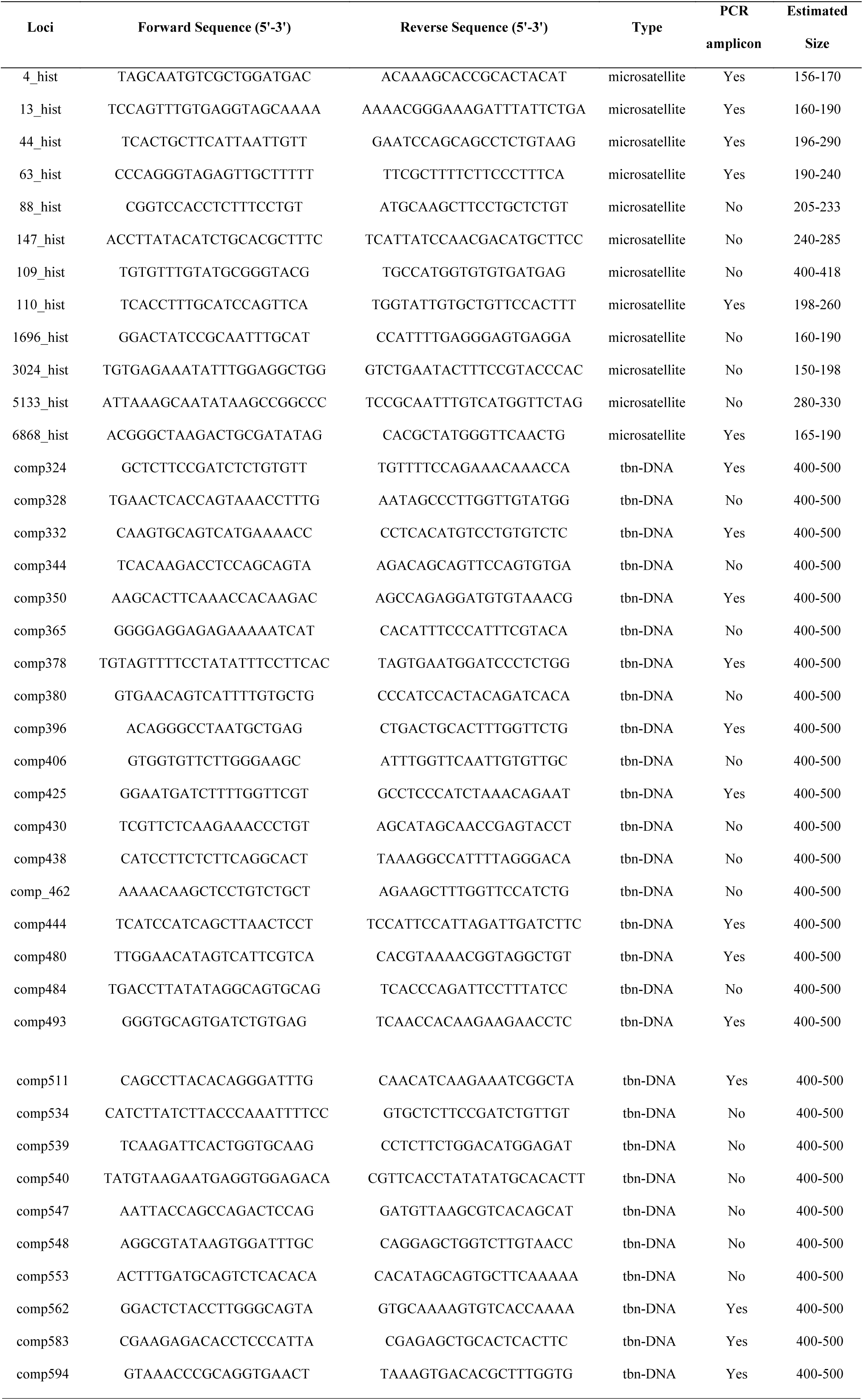
Description of nuclear markers implemented in this study.

**Table S3.** Quality descriptors of the assembled transcriptome used for nuclear loci markers design.

| Species | *O. lehmanni* |
| --- | --- |
| **File size (Mb)** | 77.3 |
| **# sequences (Contigs)** | 117,082 |
| **Total bases (b)** | 66,367,430 |
| **Mean sequence length (b)** | 566.8 |
| **Minimum length (b)** | 201 |
| **Maximum length (b)** | 12,906 |
| **N50 contig size (b)** | 809 |
| **N75 contig size (b)** | 353 |

## APPENDIX 2: Supplementary Methods

*Mitochondrial genetic clustering*

To determine the number of mitochondrial clusters (KMT), phylogenetic trees were constructed under different Bayesian (BI) and maximum likelihood (ML) optimality criteria. First, using MEGA6 (Tamura et al. 2013), we obtained an ML genealogy using extensive (level 5) Subtree Pruning and Regrafting (SPR) heuristic searches. The relative support for each node was estimated by generating 1,000 bootstrap replicates. Second, we constructed a BI-based tree using BEAST v. 2.1.2 (Drummond et al. 2012; Bouckaert et al. 2014) with a relaxed molecular clock and an uncorrelated log-normal (UCLN) model of molecular rate heterogeneity. We ran three chains for 10 million generations sampled every 100 steps. The resulting trees and log files of the three independent runs were combined using LOGCOMBINER 2.1.2. and resampled at a lower frequency (i.e. 400 steps or ¼) to reduce the number of final trees to 52.500 (Bouckaert et al. 2014). For each estimated parameter, convergence was assessed using TRACER 1.6 (http://tree.bio.ed.ac.uk/software/tracer/) and effective samples sizes (ESS) were calculated to ensure adequate mixing (ESS>320, after 30% burn in). We summarized the posterior probability density of the combined tree and log files as a maximum clade credibility (MCC) tree using TREEANNOTATOR 2.1.2 (Bouckaert et al. 2014). We visualized all trees using FIGTREE 1.4.0 (http://tree.bio.ed.ac.uk/software/figtree/). The phylogenetic relationship among sampling locations was plotted onto a TCS haplotype network with a connection limit of 95% (Clement et al. 2000) implemented in the program PopART (Population Analysis with reticulate trees) available at http://popart.otago.ac.nz/. Finally, to delimit independent evolving linages we performed a Bayesian implementation of the PTP model for species delimitation (Zhang et al. 2013) at the bTPT online server (http://species.h-its.org/) with MCMC generations = 1,000,000, thinning = 1000 and Burn-in = 0.1.

*Microsatellite molecular markers*

Microsatellite markers were generated as follow: genomic DNA from a single individual of the known O. histrionica nominal species was isolated as described in the Methods section of the main manuscript. The genomic sample was sequenced on one-eighth of a plate using 454GS-FLX technology (Roche Applied Science) at the Cornell University’s BioResource Center. We obtained 14,992 fragment reads with lengths ranging from 150 to 825 bases. We searched for di-, tri-and tetra-nucleotide tandem repeats from the dataset using MSATCOMMANDER (Faircloth 2008; Pino et al. 2012). The online version of PRIMER3 (Rozen and Skaletsky 2000) was used to access the suitability for annealing temperature and single primer-pair for each loci were chosen based on the highest PRIMER3 assigned score. Primer pairs were tested using conventional PCR in twelve individuals from two geographic localities and those loci that were successfully amplified (Table S3 in Appendix 1) were used to genotype 310 individuals from 15 different geographic locations (Table S1 in Appendix 1). We followed previously described protocols for multiplex PCR conditions, allele detection and allele calling (Anttila et al. 2014).

*Nuclear loci for multiplex PCR*

First, we generated a skin transcriptome by Trizol (InvitrogenTM) extraction of total RNA from skin portions of one known individual of O. lehmanni. Paired-end libraries (150-mer, x2; insert lengths ~300 bp) were synthesized using the Genomic Sample Prep kit (IlluminaTM) according to manufacturer’s instructions. Library quality was assessed using a 2100 Bioanalyzer (Agilent). Libraries were sequenced on an Illumina Hi-Seq 2000 with a paired-end module. Recognition, sorting, trimming of tags and quality control of the raw DNA-sequence reads were performed in FLEXBAR (Dodt et al. 2012). The remaining reads (~128 million) and the TRINITY platform (Haas et al. 2013) were used for the generation of a de – novo transcriptome. Further information about transcriptome composition and quality is presented in Table S4 in Appendix 1.

*Allele-frequency genetic clustering*

In order to assign individuals of the O. histrionica complex to genetic clusters, we implemented Bayesian clustering algorithms using a combination of allele frequency data from microsatellite (n=6) and transcriptome-derived haplotypes (n=13) in STRUCTURE v. 2.3.4 (Pritchard et al. 2000; Hubisz et al. 2009). A model assuming admixture and correlated allele frequencies was implemented in 10 independent runs with one million MCMC iterations (burn-in = 100000) for a series of clusters (KNUC) ranging from 1 (panmixia) to 15 (maximum number of localities sampled). Then, we used the statistic ΔK to select the value of K with the uppermost hierarchical level of population structure in our data (Evanno et al. 2005). For detecting species clusters and individual assignment while treating KNUC as a random variable, we also implemented STRUCTURAMA v. 1.0 (Huelsenbeck et al. 2011). The models were run for 10 million generations, sampling every 100th cycle and a burn-in period of 10000 samples. A different set of prior values was used for the shape and scale of the gamma distribution of KNUC (shape: scale = 2.5:0.5, 2.5:1, 5:2, 10:1) (Edwards and Knowles 2014). We chose the KNUC with the highest probability and individuals were assigned to species cluster by using the posterior probability of cluster assignment. Finally, for the multivariate method the combined allele frequency data was converted to synthetic variables by PCoA of a Nei’s genetic distance matrix as implemented in GenAlex v. 6.5 (Peakall and Smouse 2012) and clusters were visually inspected from PCoA dispersion plots.

## APPENDIX 3: Supplementary Figures

**Figure S1.**
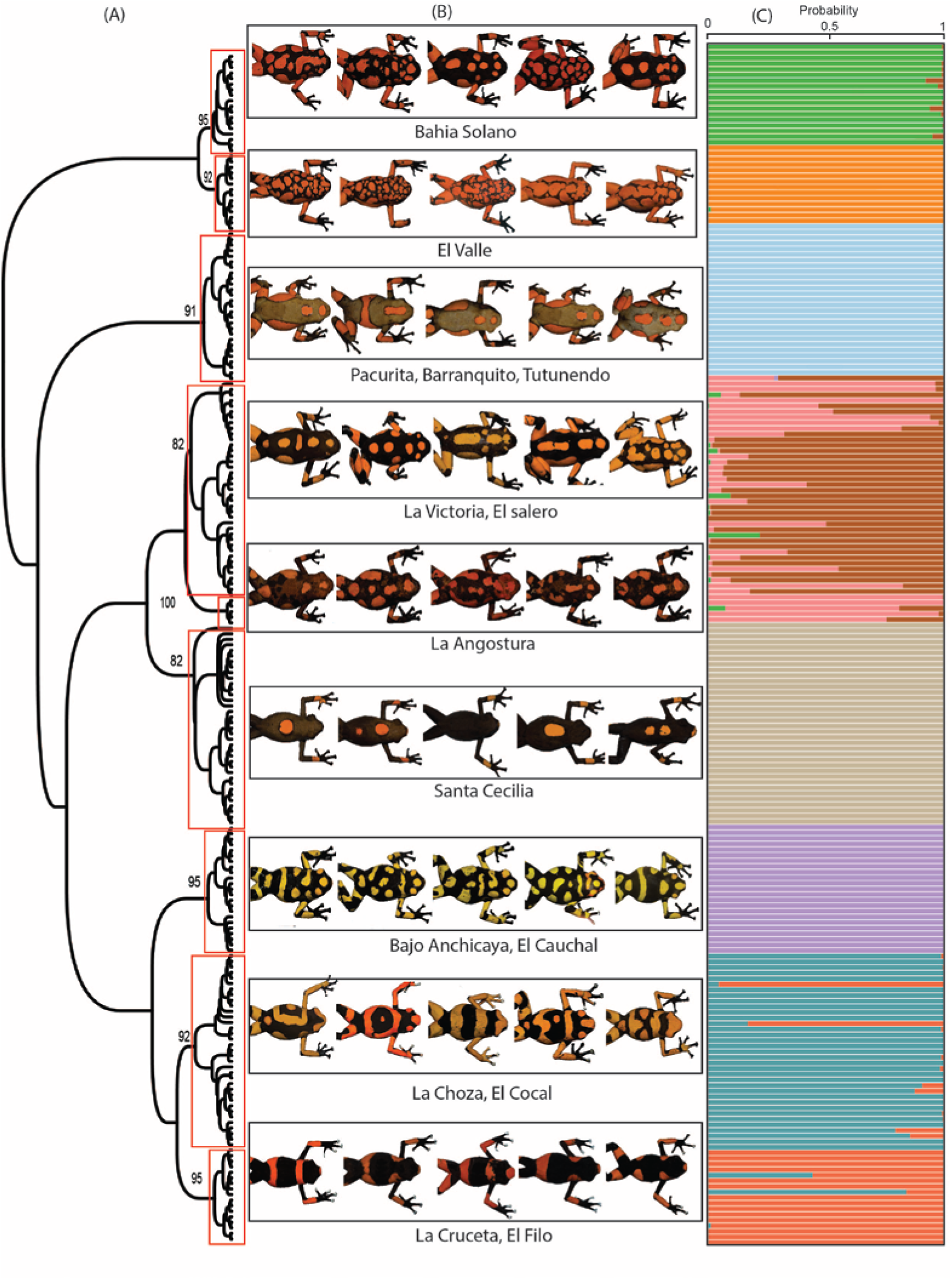
a) Morphological hierarchical clustering of individuals of the *O. histrionica* complex. Clusters with node support >82% are shown in red boxes. b) Individual morphologic variation within clusters and their respective sampling location. c) Posterior assignment probabilities (PP) for each individual based on DAPC analysis of 50 morphologic variables.

**Figure S2.**
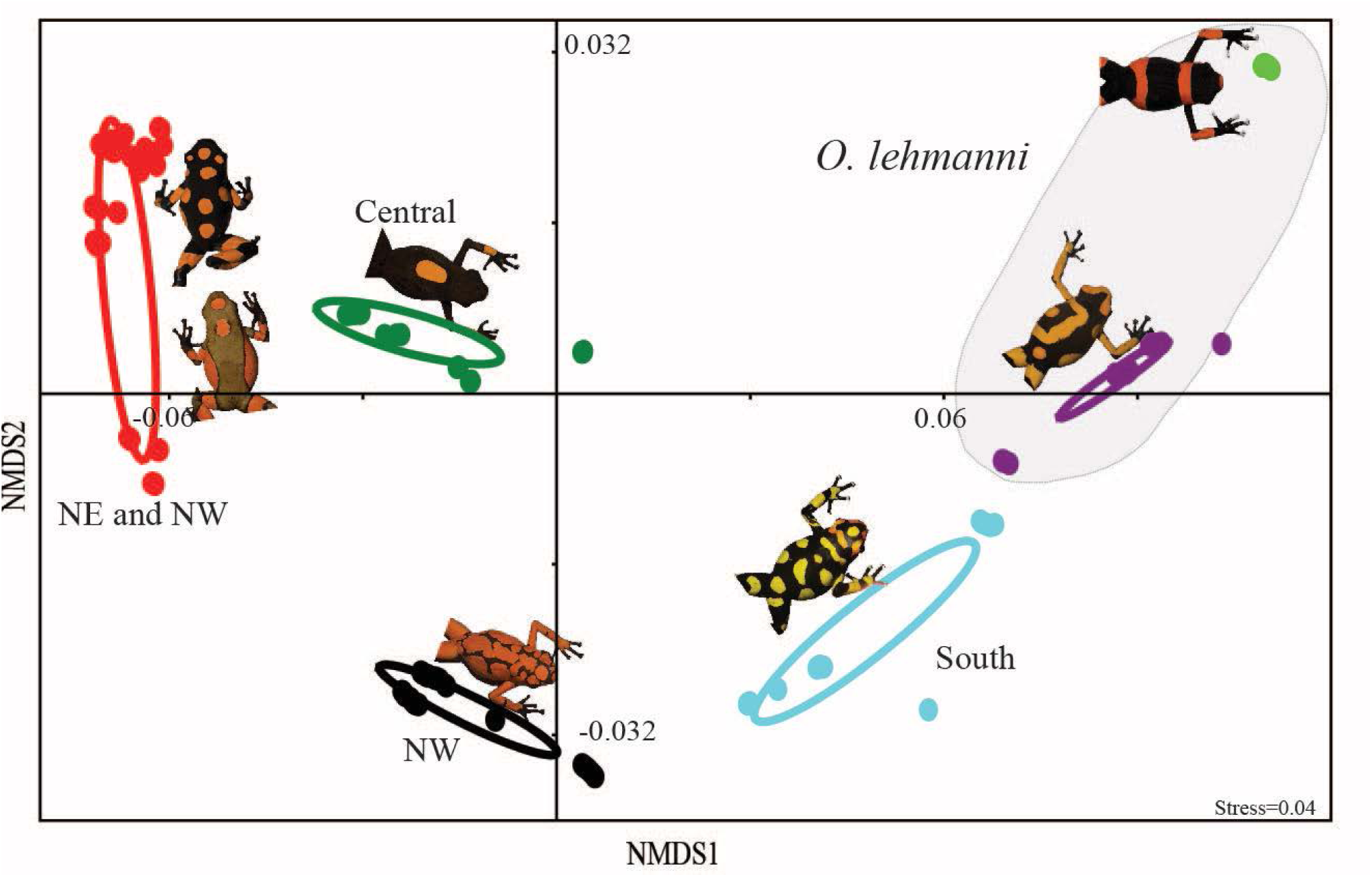
Environmental clusters by NMDS of climate data. Six potential climatic niches were found that correlates with geographic distribution. Clusters within coloured ellipses were significantly different (P<0.0001 after Bonferroni correction) and include 95% of the data. Clusters within the shadowed ellipse correspond to *O. lehmanni* populations.

**Figure S3.**
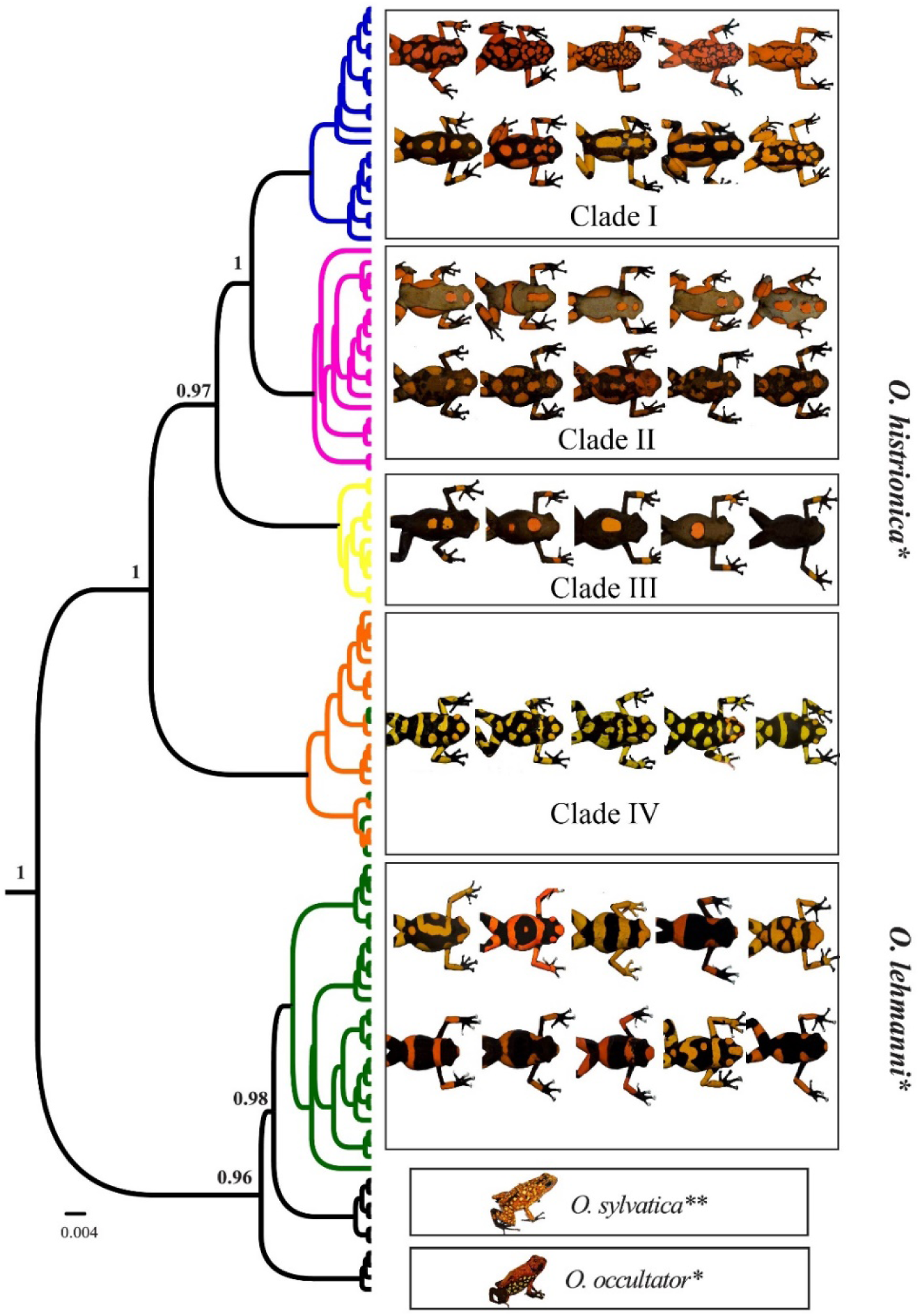
Mitochondrial species delimitation of the *O. histrionica* species complex. Numbers in nodes represent Bayesian posterior probabilities (speciation probability, SP) and only values higher than 0.95 are shown. The currently recognized *O. sylvatica* and *O. occultator* nominal species are included as control species. Species names in the figure correspond to the current taxonomic classification (*: *sensu lato* Myers and Daly 1976. **: Funkhouser 1956).

**Figure S4.**
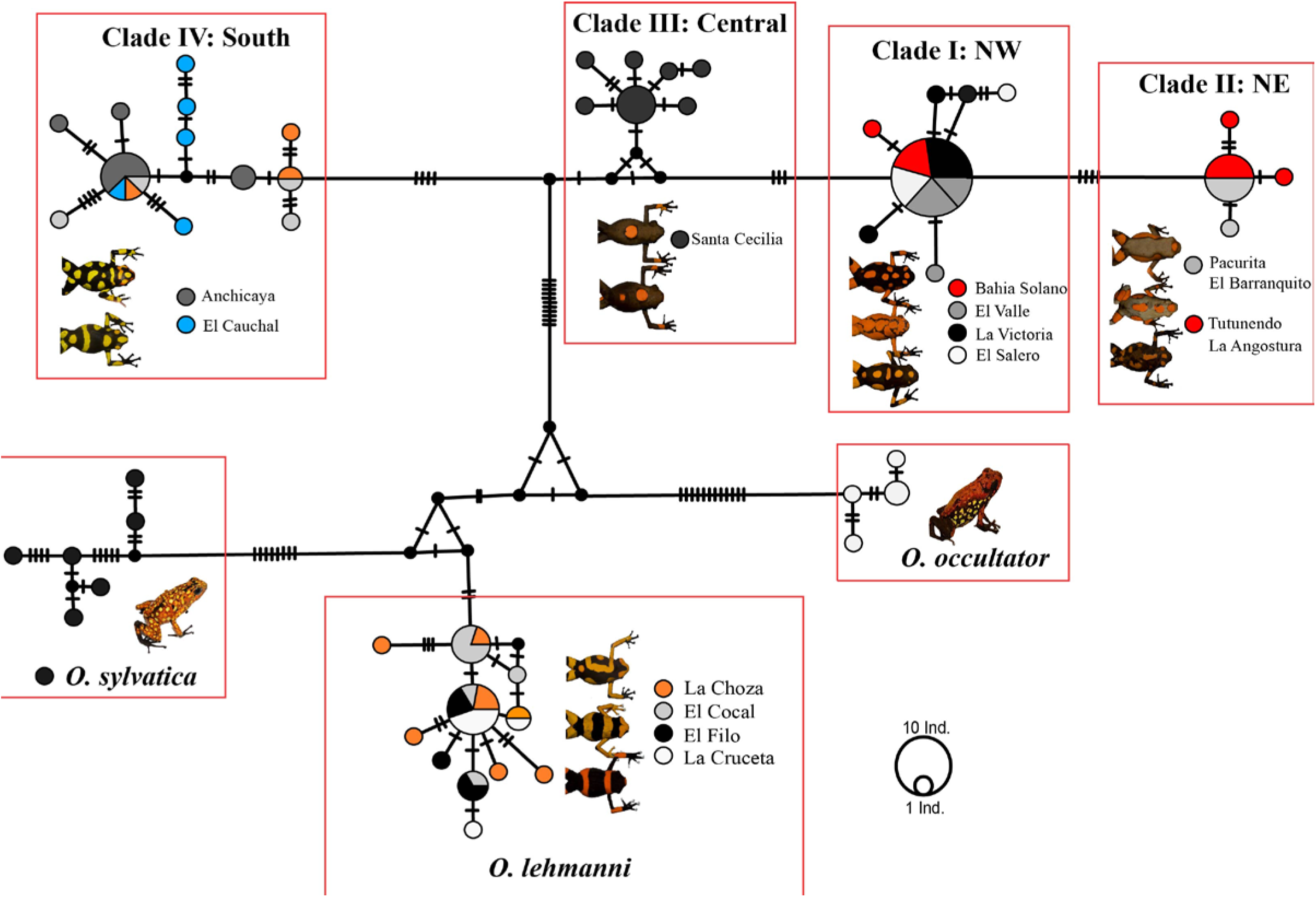
Mitochondrial (COI) parsimony haplotype network of individuals of the *O. histrionica* species complex. Clade I-IV correspond to those mitochondrial clades of Figure S3 in Appendix 3. *O. sylvatica* and *O. occultator* are included as control species.

**Figure S5.**
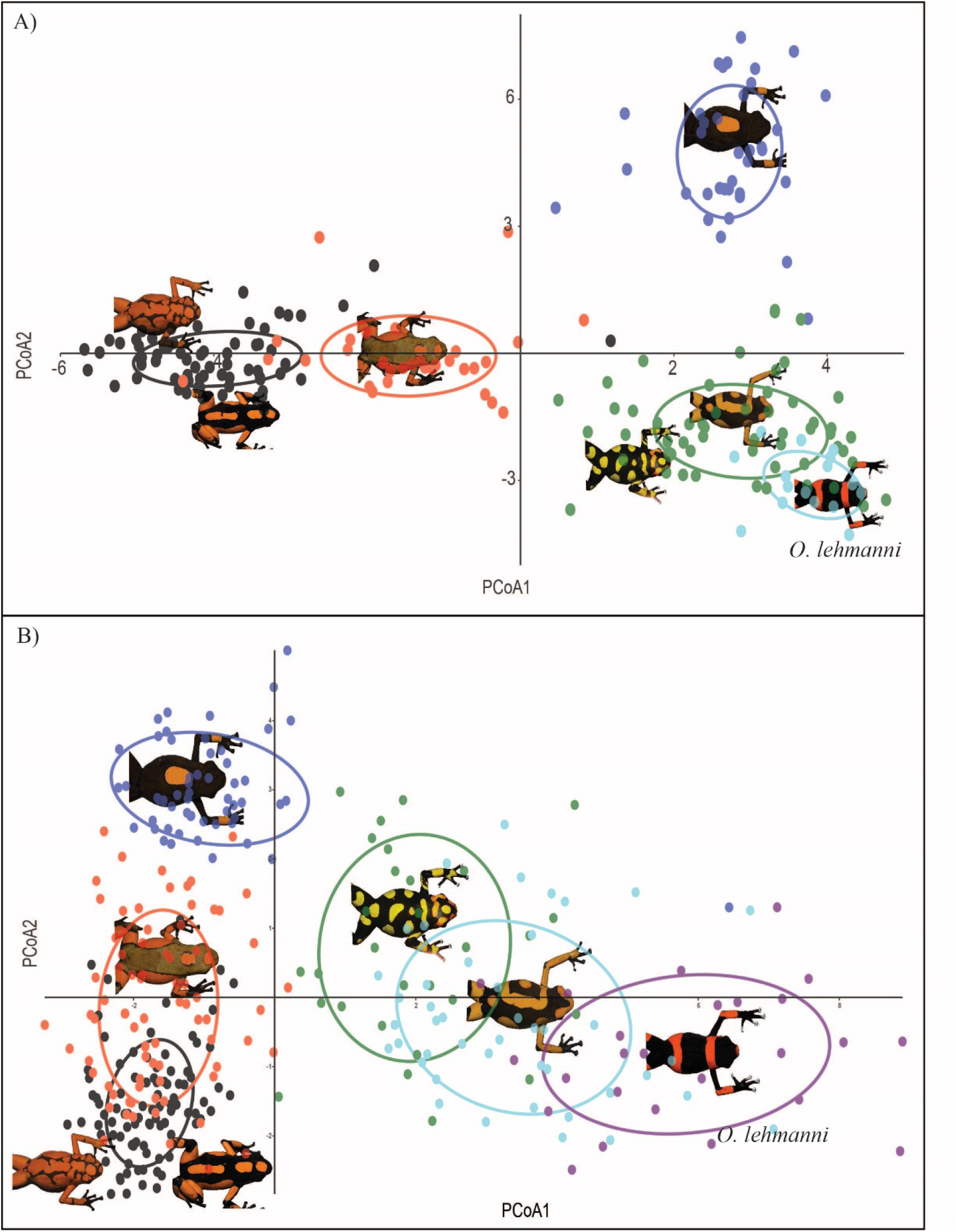
Genetic clusters by PCoA of allele frequency data from microsatellites (a) and transcriptome-based markers (b). Five cluster were detected by microsatellites markers while next generation sequencing data suggested the formation of six distinct clusters.

**Figure S6.**
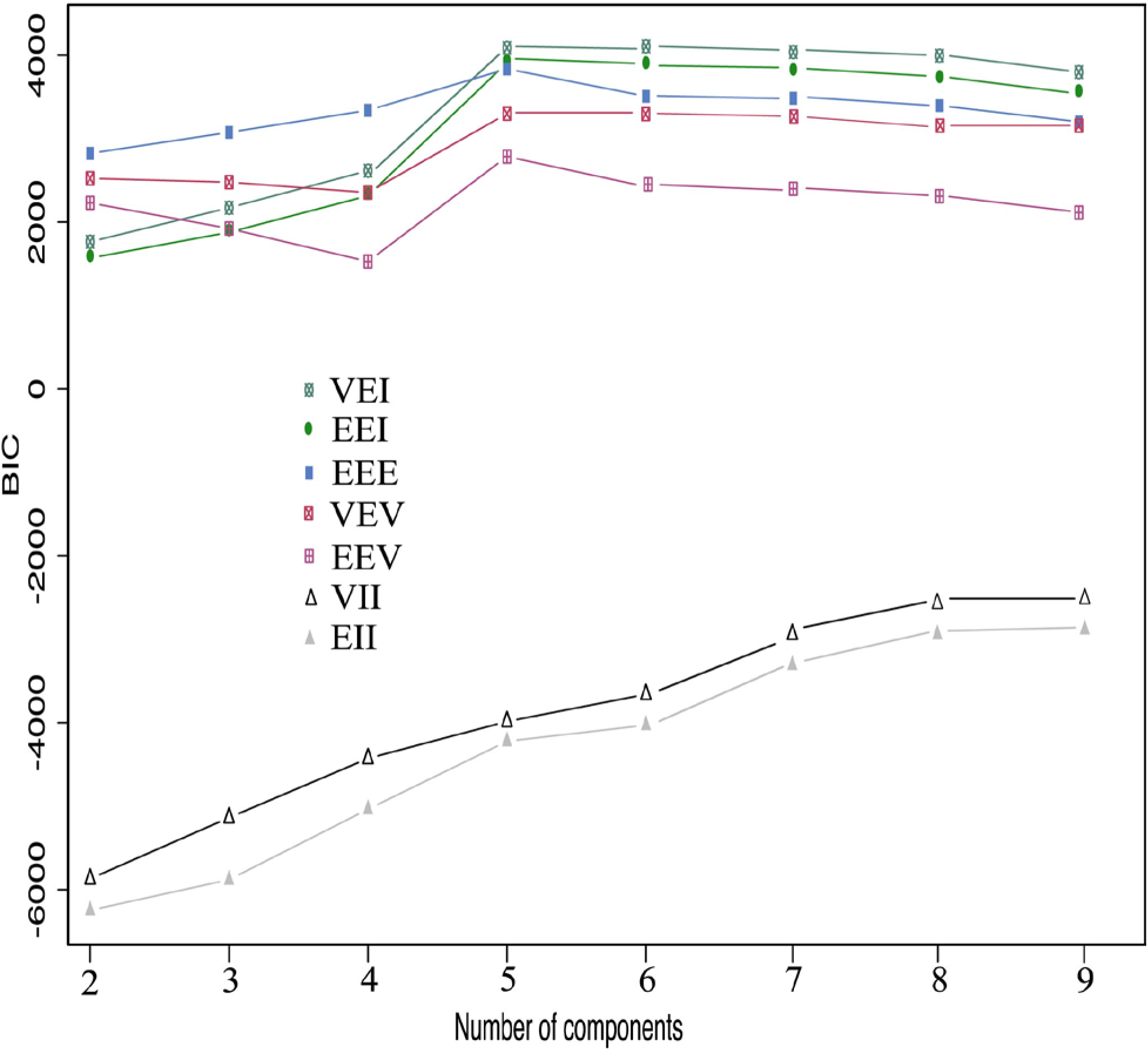
Gaussian clustering graph showing a plateau in BIC value at K=5. According to the *Mclust* analysis, the best model corresponded to an equal-covariance and diagonal distribution model (VEI) with five components or clusters.

